# Unsupervised Representation Learning of C. elegans Poses and Behavior Sequences From Microscope Video Recordings

**DOI:** 10.1101/2025.02.14.638285

**Authors:** Maurice Deserno, Katarzyna Bozek

## Abstract

Caenorhabditis elegans (*C. elegans*) is an important model system for studying molecular mechanisms in disease and aging. The nematode can be imaged in highly parallel phenotypic screens resulting in large volumes of video data of the moving worm. However converting the rich, pixel-encoded phenotypical information into meaningful, quantitative description of behavior is a challenging task. There is a range of methods for quantification of the simple body shape of *C. elegans* and the features of its motion. These methods however are often multi-step and fail in the case of highly coiled and self-overlapping worms. Motivated by the recent development of self-supervised deep learning methods in computer vision and natural language processing, we propose an unbiased, label-free approach to quantify worm pose and motion from video data directly. We represent worm posture and behavior as embedding vectors and visualize them in a unified embeddings space. We observe that the vector embeddings capture meaningful features describing worm shape and motion, such as the degree of body bend or the speed of movement. Importantly, using pixel values directly as input, our method captures coiled worm behaviors which are inaccessible to methods based on keypoint tracking or skeletonization. While our work focuses on *C. elegans*, the ability to quantify behavior directly from video data opens possibilities to study organisms without rigid skeletons whose behavior is difficult to quantify using keypoint-based approaches.

## 1 Introduction

Behavior is a window to an animal’s nervous system. Precise quantification of behavior allows to determine fine phenotypic effects of genetic mutations or pharmacological interventions and, eventually, their underlying neural mechanisms. Keypoint tracking methods and motion tracking imaging systems have enabled acquiring precise information on animal posture and its change in time in natural settings^1–3^. It is however unclear how to quantitatively measure behavior of invertebrate species with flexible bodies and appendages. Organisms such as worms lack natural skeletons and hence distinct keypoints on their bodies. The shape of *C. elegans* is typically represented as its central body line and reduced to eigenworms^4^ that enable quantification e.g. of the motion features and dynamics. However, this approach fails in the case of coiled or self-intersecting poses of *C. elegans* and current solutions apply multi-step approaches^5^ to resolve these shapes.

Here we present a method for quantification of *C. elegans* motion based on video recordings directly. Unlike keypoint- or central body line-based approaches our method does not estimate the body structure but quantifies the behavior from the raw pixel values. Our method does not require any annotations but relies on self-supervised learning approach to learn sequence representations. This combination of self-supervision and keypoint-free pose estimation enables to forgo skeletonization and feature engineering which allows studying the full repertoire of *C. elegans* poses and behavior in a comprehensive manner.

## 2 Related Work

Previous tracking and pose estimation methods for *C. elegans* enabled a quantitative, automated analysis and a better understanding of its poses and behavior^4,6–11^. These methods allowed to comprehensively analyze worm behavior and to better understand phenotypic effects of genetic mutations, disease, or aging.

Stephens et al.^4^ tracked *C. elegans* in microscopy videos and approximated their pose with a curve. They found that approximately 95% of the total variance in angles along the curve is represented by four eigenvalues. Based on these findings they introduced the term *eigenworms* as “templates” to describe the *C. elegans* poses. Javer et al.^9^ developed a widely used single- and multiworm tracking software called Tierpsy. The software segments *C. elegans* and estimates their outlines and the skeleton. Additionally, it computes several hand-engineered features characterizing pose and motion of an individual e.g. the 6 eigenworms^4^, the maximum amplitude of the skeleton’s major axis, the degree of bend of different body segments, different body size measurements (such as length and width) or the motion mode (backward or forward). Both the eigenworms quantification and Tierpsy are based on classical computer vision approaches and do not allow to quantify coiled or overlapping poses of the worm. These poses are inaccessible to these methods.

Several methods address the challenge of accurate estimation of coiled and (self-)intersecting poses. WormPose^5^ is a Residual Network^12^(ResNet)-based method applying a multi-step approach that allows for estimating poses of coiling worms. To estimate the center line using equidistant keypoints, the method relies on video data with detected/annotated center lines (e.g. by Tierpsy) for frames prior to the occurrence of coiling behavior. The authors train their network with synthetically generated images of *C. elegans* to avoid time-consuming human labeling. The network learns to predict the two different centerlines resulting from different head/tail orientations. During evaluation a synthetic image is generated for each predicted centerline. By comparing the generated images to the input the best prediction is determined. Recent methods like DeepTangle by Alonso and Kirkegaard^13^ and its extension DeepTangleCrawl^14^ by Weheliye et al. enable robust skeletonization and tracking of *C. elegans* with overlaps and on a noisy background and allow better phenotypic screening. Still these methods fail when *C. elegans* are very tightly coiled or individuals lie parallel to each other over an extended time.

## 3 Methods

### 3.1 Datasets

All data we used for our experiments are publicly available on Zenodo^15^ in the *Open Worm Movement Database*^9^. The data was downloaded using a python script filtering for specific parameters such as strain. For accessing the repository, we used the Open Archives Initiative Protocol for Metadata Harvesting (OAI-PMH). We created two datasets grouping genetic strains and long-term recordings of single individuals, respectively. The first dataset consists of seven genetic strains with one of them being the wild type. In the following we refer to this set as *strain dataset*.

This dataset includes a total of 165 videos of the following strains:

- N2
- AQ2932 (nca-2(gk5)III; unc-77(gk9)IV; nzIs29[punc-17::rho-1(G14V); punc-122::GFP])
- AQ2934 (nca-2(gk5);nzIs29[punc-17::rho-1(G14V); punc-122::GFP])
- TQ225 (trp-1(sy690)III)
- DG1856 (goa-1(sa734)I)
- DA609 (npr-1(ad609)X)
- VC731 (unc-63(ok1075)I)
- CB1141 (cat-4(e1141)V)

The second set consists of 71 videos of three strains with the same individuals recorded every day for multiple days (between 15 to 24 days per individual) during their adulthood. We call this last set the *aging dataset*. This dataset includes following genetic strains:

- AQ2947 (CGC N2 (Bristol, UK))
- OW940 (zgIs128[P(dat-1)::alpha-Synuclein::YFP])
- OW956 (zgIs144[P(dat-1)::YFP])

The data consists of video frames with masked background. Video data was recorded with frame rates varying between 25 frames per second (fps) and 32 fps. The videos of both sets have a length of almost 15 minutes each. Using Tierpsy^9,16^ we calculated features of poses and motion in all our recordings. These features together with the worm genetic strain and age represent the metadata we use for interpretation of the image and sequence representations we developed in this study.

**Figure 1.**
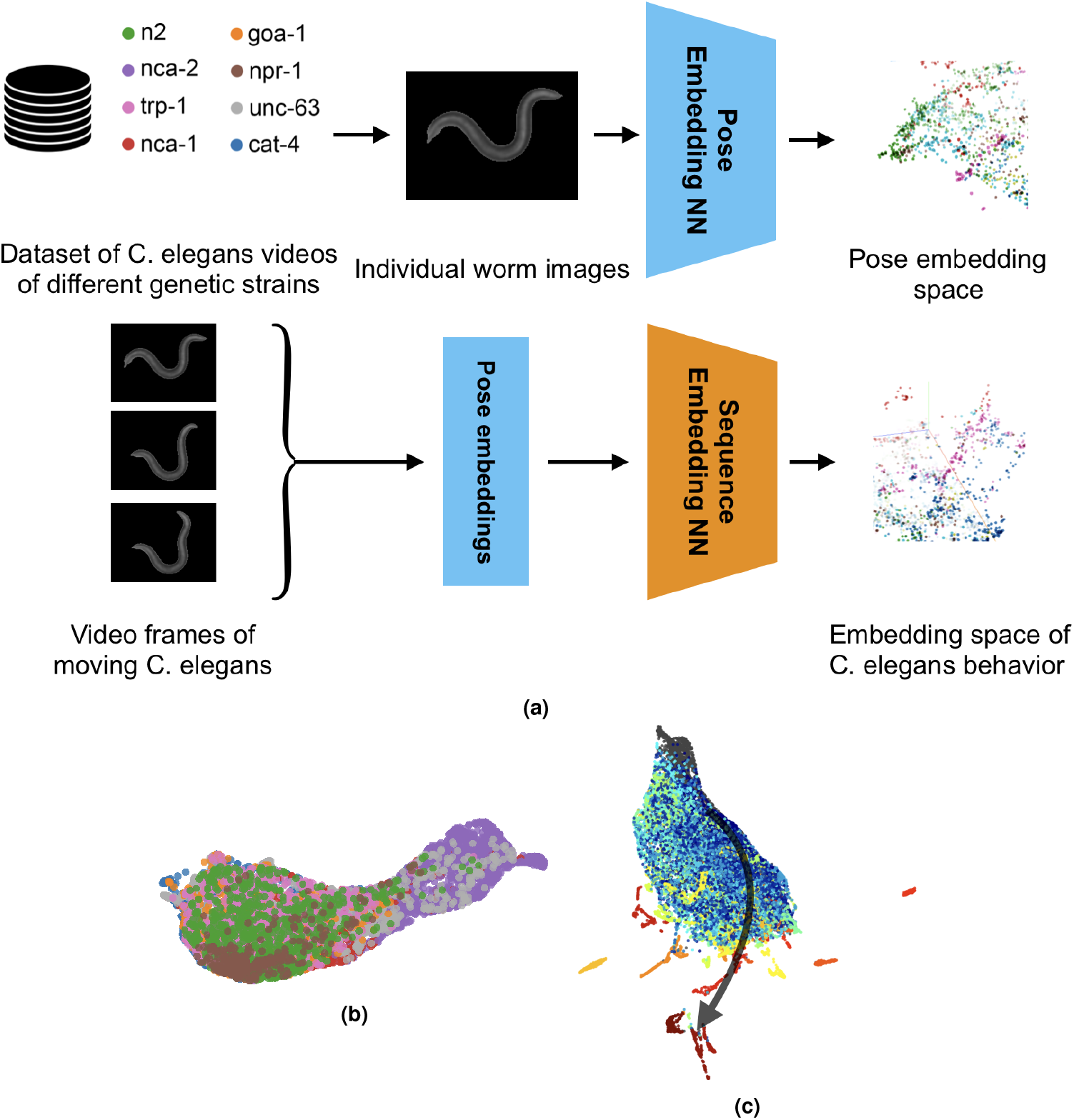
Processing pipeline and behavior representation. (a) Processing pipeline overview. We use a large set of video data of worm genetic strains and employ a contrastive learning approach to encode individual poses of the worm directly from the video frames. We next inspect these pose embeddings using their visualization in a 3D scatter plot. The trained pose embedding network is used to embed each video frame which is next an input to the sequence embedding network. Similarly to pose embeddings, we inspect the embedding space of worm behaviors using visualization techniques and motion features quantified with Tierpsy. (b) Visualization of the strain dataset behavior embedding space colored by the underlying genetic strain. (c) Visualization of the aging dataset behavior embedding space, illustrating the behavioral change with age in the direction of the arrow moving from young (blue) to old (red).

### 3.2 Data pre-processing

The background of the *C. elegans* images downloaded from Zenodo^15^ is masked with black pixels. In some images the masking contains errors with background objects not masked out. Here we apply a combination of different methods including connected components and morphological operations to filter out smaller foreground blobs and to better match the background mask to the worm shape (similar to^5^). As a result we remove the errors and limit the foreground to one object only - the worm (see Fig. 2 “Data Preprocessing”). Further, we change the background mask pixel value from black (0) to gray (127). Next, we crop the foreground in the image, pad and resize it to a common image size of 128 × 128 pixels. This way, we remove excessive background pixels and center the object in the middle of the image while preserving its relative size. In the final step, we apply Principal Component Analysis (PCA) to rotate the object to a vertical orientation (Fig. 2).

**Figure 2.**
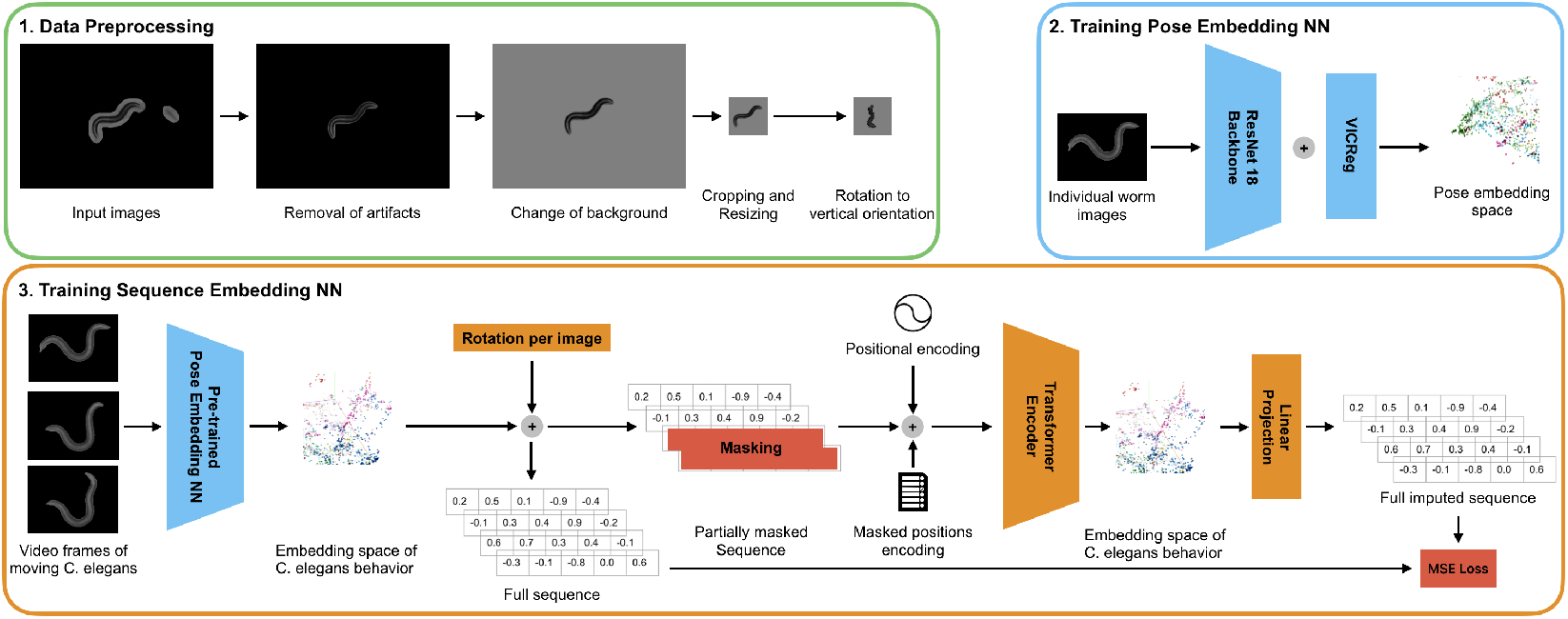
Data preprocessing and network architecture. 1) Data preprocessing pipeline: Artifacts are removed keeping only the worm as foreground object. We change the background to gray, crop the image to keep the worm centered and resize it to 128 × 128*px*. Finally, we rotate the worm to a vertical orientation. 2) A contrastive learning network is trained with images in random order to learn pose embeddings. 3) Using the ResNet-18 trained in (2) we embed sequences of 12 frames of moving *C. elegans*. Rotation information is concatenated with the encoded sequences and the last 5 frame embeddings are masked out. A Transformer-encoder learns behavior embeddings by imputing the masked sequence elements.

We store the degree of rotation in addition to the strain and the day of adulthood in the aging dataset as the metadata. The metadata is not used in model training but in the model interpretation and visualization. We split the dataset into train, validation and test subsets with the proportions 0.76, 0.10, 0.14 and save the video frames as PyTorch^17^ tensors in a .pt file per subset.

Following the data pre-processing, we train our deep learning approach, which includes two parts. The first part consists of a contrastive learning method to represent spatial poses of *C. elegans* based on their images. The second part is a Transformer encoder architecture that uses the learned pose representations to predict masked parts of a spatiotemporal sequence. In the following, we describe the two parts in detail.

### 3.3 Contrastive Learning for pose representations

We apply contrastive learning to learn representations of poses from *C. elegans* image data. It is a self-supervised approach that does not require labels. Specifically, we use a version of VICReg^18^ adapted to our task. As backbone we chose ResNet18 over ResNet50 originally used in VICReg because of its smaller size. Our experiments suggested that the results do not improve using a larger feature extractor. We use a modified set of augmentations to ensure the network focuses on the important pose differences and learns to embed them rather than embedding the differences in e.g. lightning conditions or size of individuals. The output dimensionality was set to 64 with a hidden network dimensionality of 128.

To avoid having many similar poses in the training set, we subsample video frames by a factor of 10. We train the network using a batch size of 512 for 80 epochs on a NVIDIA Tesla V100-SXM2 with 32 GB of memory. As optimizer we chose AdamW^19^ with a learning rate of 0.001. Additionally we use Cosine Annealing^20^ as learning rate scheduler. The loss is calculated the same way as proposed by the authors of VICReg^18^: a weighted combination (compare with 1) of variance *v*, invariance *s* and covariance *c* loss with weights set to *µ* = 25.0, *λ* = 25.0 and *ν* = 1.0.

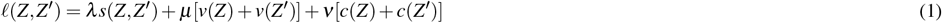

### 3.4 Transformer encoder for sequence data imputation

To integrate the temporal component of behavior into the learned embeddings we employ a Transformer encoder neural network architecture^21–23^. The Transformer encoder consists of a multi-head attention block and a feed-forward network. This type of architecture has been used primarily in natural language processing (NLP) (e.g. by BERT^22^) and was later adapted to images (e.g. in Vision Transformer^23^). We attach the pre-trained pose representation network (see 3.3) as backbone to the Transformer network and freeze this backbone. We add a linear projection network to the last layer of the Transformer encoder network that infers embeddings of individual poses in the sequence.

During training, we input 12 ordered video frames as a sequence into the pre-trained pose representation network to generate pose embeddings. Here, we downsample the videos by a factor of 5 which is sufficient to capture the worm’s motion in a smooth manner. With frame rates between ∼25 − 32 fps (see section 3.1) this results in a sequence covering between ∼2 - 1.6 seconds in real time. We store the ordered pose embeddings generated by the pose backbone as ground truth information for later evaluation.

Next, we construct sequences of 12 consecutive frame embeddings and attach frame rotation information generated during pre-processing. We mask the last 5 sequence elements by replacing them with zeroes (similar to^22,24^) before passing the sequence to the transformer network. We add sine-cosine positional encoding^21^ and masked position encoding to the pose embeddings. The masked position encoding is a vector, indicating if a sequence element (frame) is masked (value 1 in the vector) or is not masked (value 0 in the vector)^24^. This vector is embedded and then added the same way as the positional encoding^25^ (see Fig. 2). The pose embeddings together with positional and mask embeddings are the input to the transformer encoder. The Transformer network is trained to impute the missing values in the sequence. Using the linear projection network, pose embeddings and their rotations are predicted for each of the masked positions. We calculate the Mean Squared Error (MSE) loss between the embeddings generated by the pose representation network and the predictions of the linear projection network.

Pose representations have a dimensionality of 64 (see 3.3). The transformer uses a hidden dimensionality of 128 and consists of one encoding block and two heads. For training we use AdamW^19^ as optimizer with a learning rate of 0.0005 for 250 epochs with a batch size of 64. The network was trained and tested on a NVIDIA Tesla V100-SXM2 with 32 GB of memory.

### 3.5 Visualization of pose and motion embeddings

To inspect the *C. elegans* pose and sequence embeddings we use the dimensionality reduction technique Uniform Manifold Approximation and Projection (UMAP)^26^. By applying UMAP, we reduce the embeddings to three dimensions to visualize them as scatter plots. We used the python implementation^1^ of UMAP with the parameters *n_neighbors=30, min_dist=0*.*25, n_components=3* and *random_state=42* for the pose embedding space and for the behavior sequence embedding space.

## 4 Results

### 4.1 Pose representations

We first inspected the embeddings of individual worm poses. We project the embeddings in 3D using UMAP and inspect whether the embedding space reflects Tierpsy-based^9,16^ pose features, as well as worm genetic strain. Figure 3a illustrates the pose embedding space of the strain datasets (see 3.1). This space shows a clear spatial ordering of poses according to their degree of bending (see Fig. 3a). While one end of the point cloud consists of strongly coiled worms, the opposite end clusters worms with poses close to a straight line. The points are colored according to the maximum amplitude of the bend along the worm body line. The straight poses have a low amplitude value, the more bent ones a higher one. There is a clear gradient of this value along the point cloud. However, the coiled worm shapes are missing this feature value (marked in gray color in Fig. 3a) as Tierpsy cannot resolve these poses^9,16^. This reveals an advantage of our approach: it allows to capture all worm poses, from straight to strongly coiled ones, in a uniform and smooth embedding space. Our approach groups coiling and bending poses together with a clear transition between them, whereas an important fraction of worm poses is not possible to quantify with skeletonization-based methods. Since our approach does not require any skeleton or keypoint estimations, it is robust against coiling and self-intersecting postures. A large number of embeddings in the gray area in Fig. 3a belongs to *C. elegans* of the AQ2934 strain.

**Figure 3.**
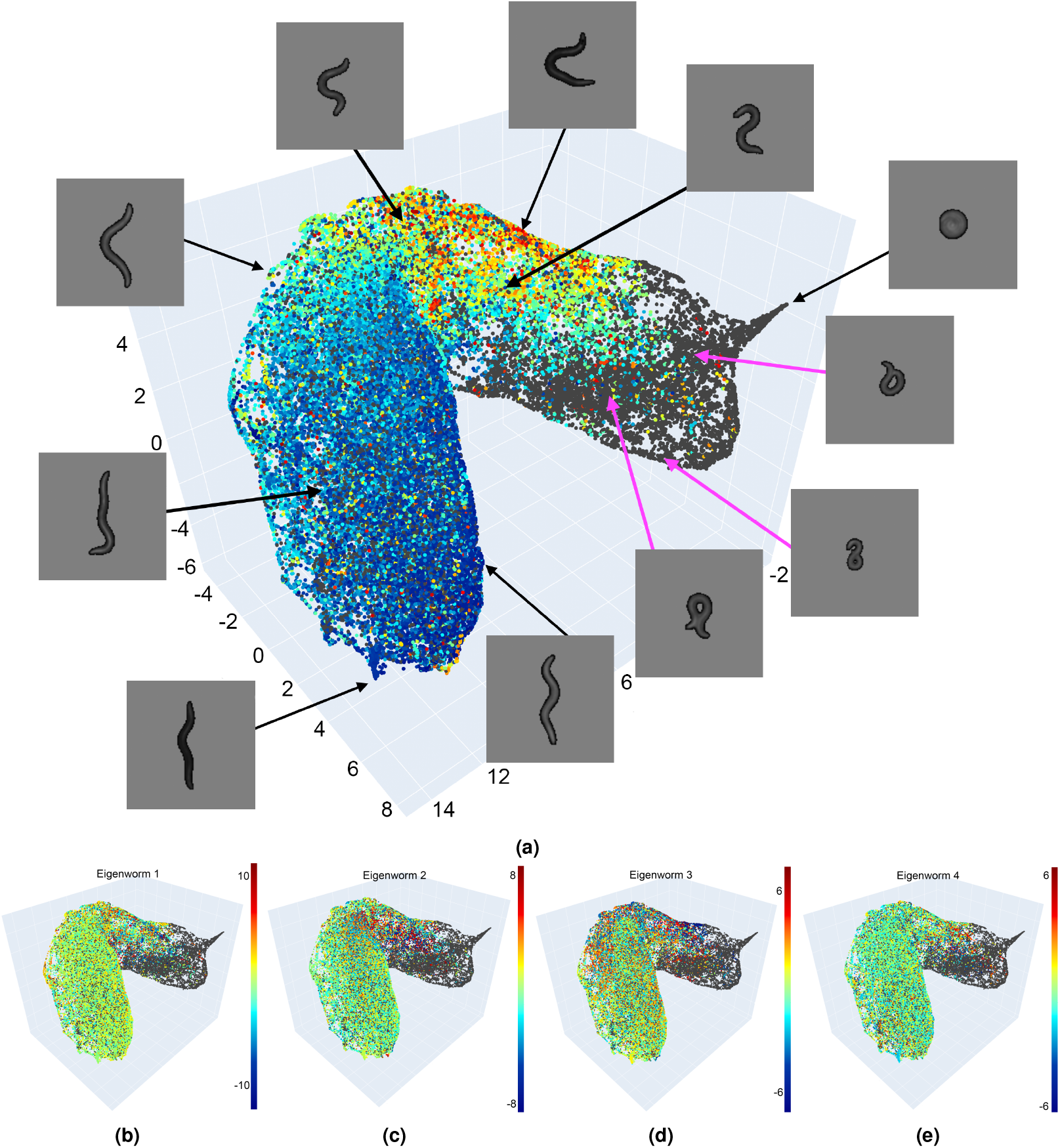
Visualization of the pose embedding space. (a) We reduced the embedding space to 3D using UMAP and colored it with the Tierpsy *max_amplitude* feature. Dark gray dots indicate poses for which this feature could not be quantified using Tierpsy. There is a gradient in coloring suggesting that similar poses occupy neighboring parts of the embedding space. Example images of poses are shown with an indication of their position in the embedding space. Strongly coiled and almost straight worms occupy opposite ends of the point cloud. (b-e) Pose embedding space colored according to their eigenworm 1 to 4 values.

### 4.2 Sequence representations

We next trained a Transformer-based approach to embed sequences of worm postures captured in a video recording. The Transformer network takes as input sequences of pose embeddings where the second half of each sequence is masked. The network is trained to infer the masked part of the sequence as well as the rotation angle of the worm in the video. The MSE of the masked pose estimation in the strain dataset is 0.106 while of the rotation angle 0.0129, which represents an error of ∼ 20.47°. Via this self-supervised approach the network learns representations of the sequences that encode worm posture, its change in time, and the dynamics of this motion in a comprehensive manner. Similar to the pose embeddings, we visualize the embedding space of *C. elegans* short-term behaviors in 3D (Fig. 4a).

**Figure 4.**
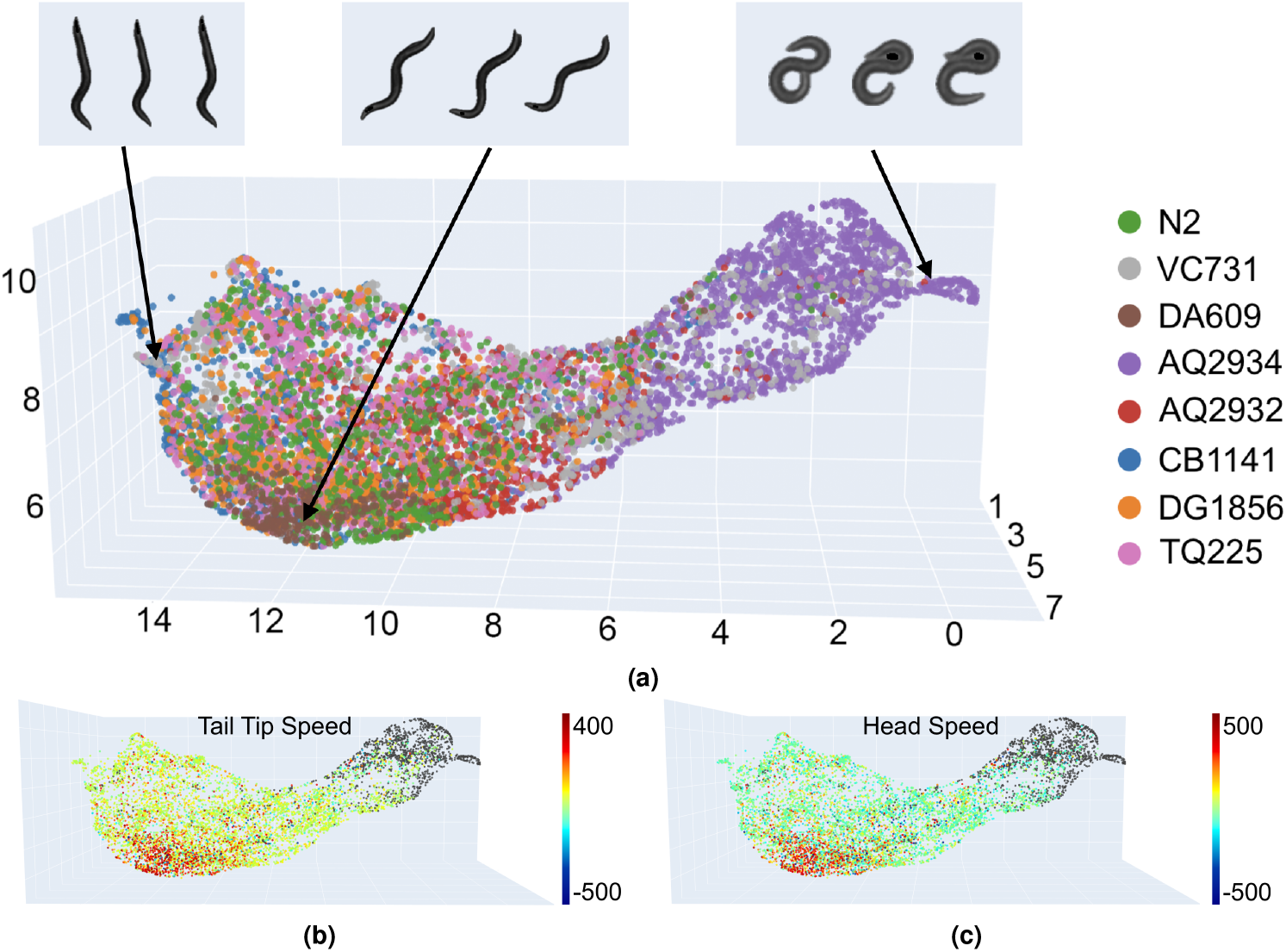
Behavior embedding space of the strain dataset. (a) Embedding space colored by strain. Worm images above correspond to 1^st^, 6^th^ and 12^th^ frame of three example sequences. (b) Embeddings space colored by tail tip speed and (c) head speed. Gray dots in (b) and (c) indicate sequences for which these Tierpsy features are missing.

This visualization shows a clear separation of sequences of the strain AQ2934 (labeled in orange) from sequences of the other strains. This separation was also present in the pose embedding space, and reflects the frequent and heavy coiling behavior of the AQ2934 strain. Behavior sequences of strain DA609 (marked in brown) are also grouped together in the embedding space. This strain is known for aggregating and burrowing behavior^27,28^. Next to the DA609 cluster is a larger area where behavior sequences of different strains mix. This likely occurs since most strains share common behaviors such as simple forward locomotion.

To further interpret the behavior embedding space, we colored it according to motion speed features quantified with Tierpsy (Fig. 4ab-c). We observed that sequences with faster movement are more frequent in the center of mass of the embedding space. This confirms our observation that crawling behavior, common to most of the strains, is located in this part of the embedding space.

### 4.3 Worm behavior changes with age

We next inspected the behavior embedding space of the aging dataset. This dataset contains 71 individuals that were recorded over their adulthood, for time span of up to 24 days. We employed our approach to inspect which parts of the embedding space those individuals occupy as they age (Fig. 5a). Young individuals appear to display a wide range of behaviors, while as they age their behavior repertoire reduces. Markedly, the patterns of aging in behavior are consistent among individuals. This can be seen in the embedding space labeled by day of adulthood (see Fig. 5a). Behaviors of individuals from day 1 to 10 span a wide area in the space, while embeddings for day 10 to 15 cover much more limited areas at the bottom part of the point cloud. From day 15 onward the embeddings almost only form outlying groupings. The reason for these behaviors to localize on the outside of the embedding space can be two-fold. On the one hand these individuals move slower and assume fewer different poses which differ from those of more agile younger individuals with their typical crawling/swimming locomotion and coiling behaviors. On the other hand, old individuals are not included in the strain dataset on which the Transformer was trained. Although the strain dataset and the aging dataset were recorded in similar ways, the behaviors of older individuals were never seen by the network.

**Figure 5.**
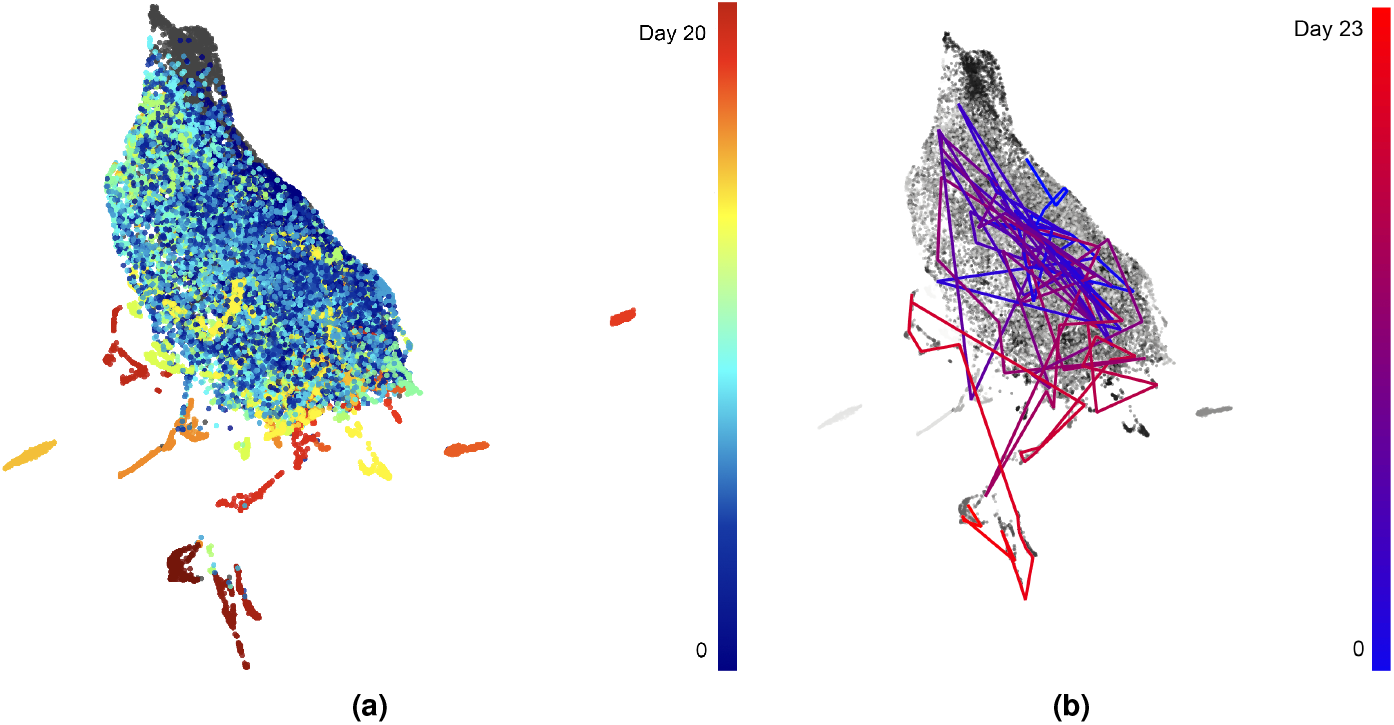
Behavior embedding space of the strain and aging datasets combined. (a) Embedding space colored by age. With age we refer to the day of adulthood of an individual *C. elegans*. Gray color indicates missing age data of worms from the strain dataset. (b) Behaviors of one individual linked over the course of its aging. Starting with blue at the last day of the L4 stage, progressing to red until the last recorded day (23) of adulthood.

Additionally to the age color-coded embedding space, we plotted the trajectory in the embedding space of one individual of strain AQ2947 (Fig. 5b). This trajectory links behaviors of this worm as it ages. It illustrates the broad variety of the behavior of this individual up to day 15 after which the its behaviors are limited to the bottom part of the embedding space.

## 5 Discussion

In this work, we presented a deep learning-based approach for representation learning of *C. elegans* poses and behavior sequences from bright-field microscopy videos without human annotations. Our method uses a combination of Contrastive Learning and a Transformer architecture originally developed for self-supervised learning in computer vision and NLP^22^. We draw inspiration from these methods to demonstrate that the pose and motion of *C. elegans* can be quantified in a meaningful manner without the use of labels. Contrary to previous approaches, our method does not require worm skeletonization, keypoints definition, or any pose or behavior categories. Our approach allows to embed all worm poses and pose sequences, from straight ones to the challenging poses of tightly coiling and strongly bending *C. elegans*. We demonstrate that, even though our methods are based exclusively on image pixel values, the resulting image and video embeddings reflect quantitative features describing the worm shape and its motion, such as degree of bend, eigenworms and speed of motion. We apply our method to the video data of different genetic strains as well as aging worms and illustrate the differences in behavior of worms of various strains and ages.

To summarize, the advantages of our approach are:

1. Embedding challenging poses without relying on annotations.
2. Quantifying previously inaccessible behaviors.
3. Capturing hand-engineered features without explicitly calculating them.
4. Ability to capture in a comprehensive manner properties of poses and behavior.

One limitation of our approach is the inability to distinguish between the head and tail of the worm. Head and tail movements are important elements of the worm behavior. Since *C. elegans* typically move head-first, the head/tail orientation can be estimated based on their direction of movement. However, for strains that frequently move backward, this rule would not apply and the head/tail orientation would need to be estimated based on their visual features. While this remains a challenging task, future work should incorporate predicted head/tail orientation as input to the network in our approach. Alternatively, video frames could be adjusted so that *C. elegans* always face head-up, rather than simply aligning all worms to a vertical orientation without considering head/tail direction.

While our approach offers many advantages over methods based on hand-engineered features, one drawback is its lower direct interpretability. For example, a feature such as head speed provides straightforward, low-level behavioral insights, whereas our embedding space visualizations combine all characteristics of the worm motion and are therefore more difficult to interpret, similar to eigenworm features^4^. On the other hand, the comprehensive motion embeddings derived from our method are a powerful representation for downstream tasks such as behavior or strain classification, reaching beyond analyses based individual motion features.

In this work we focused on behaviors spanning two seconds. Future experiments could explore embedding sequences with different time spans. Extending the length of the input video to four or eight seconds may allow to capture additional behaviors, from brief actions to prolonged activities such as mating. Longer videos can be incorporated in various ways, such as adjusting the step size between frames or increasing the sequence length.

Since pixel-based approaches like ours do not rely on skeleton or keypoints definition, they can be applied to any body form. This ability to quantify behavior directly from pixels opens possibilities to study a wide range of organisms, including cephalopods^29^ or single-celled organisms with flagella or cilia^30,31^ in a comprehensive manner.

## 6 Acknowledgments

We would like to thank Greg J. Stephens and André E. X. Brown for their valuable comments and discussions. Maurice Deserno and Katarzyna Bozek were supported by the North Rhine-Westphalia return program (311-8.03.03.02-147635), BMBF program Junior Group Consortia in Systems Medicine (01ZX1917B) and hosted by the Center for Molecular Medicine Cologne.

## 7 Author contributions statement

M.D.: methodology, software, experiments and analysis. K.B.: supervision, data and funding acquisition. M.D. and K.B. writing the article. All authors reviewed the manuscript.

## Footnotes

1 https://github.com/lmcinnes/umap

